# Modification Mechanism of Phenol-formaldehyde Resin with Crude Bio-oil by Model Compound Method

**DOI:** 10.1101/2022.07.04.498740

**Authors:** Yuxiang Yu, Xiaoqian Qiu, Chao Li, Jianmin Chang, Defu Bao

**Affiliations:** College of Art and Design, Zhejiang Sci-Tech University, 928 Seconded Avenue, Xiasha High Education Zone, Hangzhou 310018, China; Lab of Material Innovation Design and Intelligent Interaction, Zhejiang Sci-Tech University, 928 Seconded Avenue, Xiasha High Education Zone, Hangzhou 310018, China; College of Materials Science and Technology, Beijing Forestry University, 35 Qinghua East Road, Haidian District, Beijing, 100083, China

**Keywords:** Modification mechanism, Phenol-formaldehyde resin, Crude bio-oil, Model compound method

## Abstract

To clarify the modification mechanism of bio-oil for phenol-formaldehyde resin with crude bio-oil (BPF), the bio-oil compounds were simplified by model compound method according to the component distribution. The phenol-formaldehyde resin with bio-oil model compounds (BMPF) were prepared and their basic performance, bonding strength and aging characteristics were determined. The changes on the microstructure and chemical bonds of BMPF were also analyzed by scanning electron microscope, Fourier transform infrared spectroscopy, and nuclear magnetic resonance analysis. Results showed that the components of bio-oil had different influence on the performance and microstructure of BMPF, especially phenols. Structural analysis indicated that the phenols and ketones of bio-oil had positive effects on the synthesis of BMPF, while the aldehydes and acids had negative effects. But all components of bio-oil could improve the aging resistance of BMPF inordinately. These results could provide a basis for the modification of BPF.

## Introduction

Phenol-formaldehyde resin (PF) is widely used in wood-based panel because of its excellent bonding strength, thermal stability, and heat resistance, etc. [1,2]. Phenol, as the main raw material for preparing PF, however, comes from fossil sources. With the promotion of global policies on resource shortage and environmental protection, seeking or developing a renewable resource to substitute phenol for preparing PF is becoming more important. Over the past decade, researchers have successfully produced bio-based PF by replacing phenol with sustainable materials such as lignin [3-6], tannins [7,8], cardanol [9], furfural [10], glyoxal [11] and bio-oil [12-15].

Bio-oil, the main product of the rapid pyrolysis of biomass, has been successfully used in the preparation of PF resin in recent years [16-18]. Compared with petroleum-based PF, bio-oil phenol-formaldehyde resin (BPF) has similar chemical and physical properties [13,19], but a better price competitiveness [20,21]. However, the components of bio-oil are complex, including hundreds of organic components such as phenols, ketones, aldehydes, acids and sugars [17,22]. Some organic components in the bio-oil can participate in the reaction to change the structure of resin, thus affecting the properties [23]. Other components that are not involved in the reaction also have influence on the resin properties as fillers [24,25]. In order to enhance the beneficial effects and reduce the negative effects of bio-oil on resin synthesis, it is of great significance to clarify the specific reaction mechanism of the different components in bio-oil.

Model compound method is a way to use artificial compounds with same or similar structure and function to simplify the complex compound, such as bio-oil. Fortunate et al. [26] used 2-hydroxybenzaldehyde as bio-oil model compound to study the delineate effect of process variables on the catalytic hydrodeoxygenation of bio-oil by computational fluid dynamics. Tran et al. [27] studied the influence of Fe/AC and Ni/gamma-Al_2_O_3_ catalysts on the hydrodeoxygenation of woody bio-oil using guaiacol as model compound. Wang et al. [28] tried to better understand the aging mechanism of bio-oil by evaluating the aging performance for 39 kinds of bio-oil model compounds, and found that acids played an important role in the aging process, because it could be the reactant in the esterification reaction as well as the catalyst for the polymerization reaction of phenol and aldehyde.

In this study, the bio-oil was simplified into five groups, including phenols, ketones, aldehydes, acids and sugars, according to the component distribution of bio-oil by gas chromatographic-mass spectrometric (GC-MS) analysis. And the bio-oil model compounds and its component model compounds were formulated by model compound method. The phenol-formaldehyde resin with bio-oil model compounds (BMPF) were prepared and their basic performance, bonding strength and aging characteristics were determined. The changes on the microstructure and chemical bonds of BMPF were analyzed by scanning electron microscope (SEM), Fourier transform infrared spectroscopy (FT-IR), and solid state nuclear magnetic resonance (NMR) analysis.

## Materials and Methods

### Materials

Bio-oil, an acid liquid (pH 3.5), was obtained by the fast pyrolysis of Larix gmelinii (Rupr.) Kuzen in a fluidized bed at 550 °C for 2-3 s by the Lab of Fast Pyrolysis of Biomass and Productive Utilization (Beijing Forestry University, Beijing, China). And the organic components of bio-oil were displayed in Table 1 by GC-MS. Phenol, formaldehyde (aqueous solution, 37 wt.%), sodium hydroxide (NaOH), guaiacol, catechol butanone, cyclopentane dione, vanillin, furfural, D-glucose, acetic acid, phenylacetic acid were supplied by Xilong Chemical Industries, Guangdong, China. Poplar veneers (8 % moisture content, 400 mm × 400 mm × 1.5 mm) were provided by Xinda wooden Co., Ltd., Hebei, China.

**Table 1.**
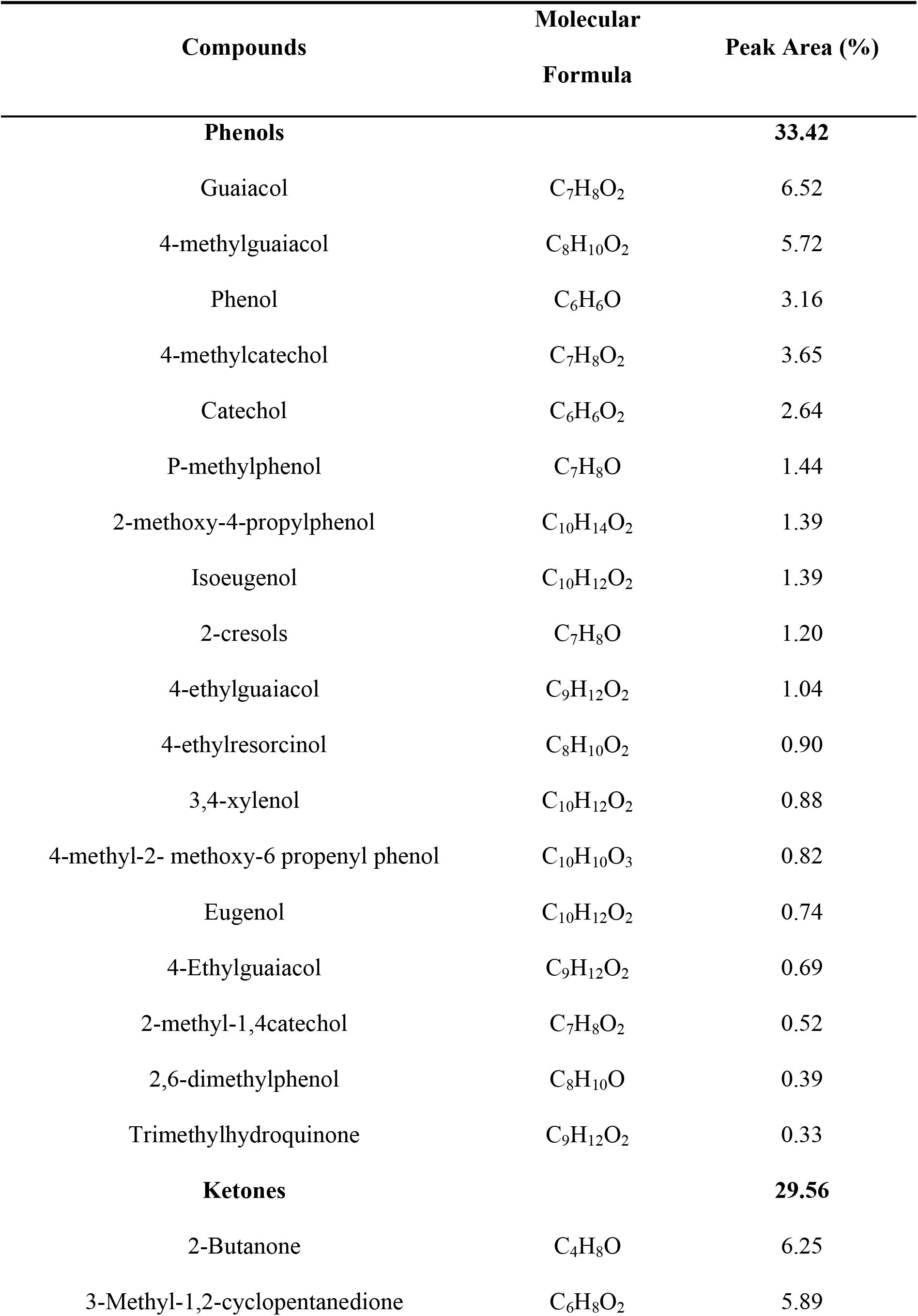
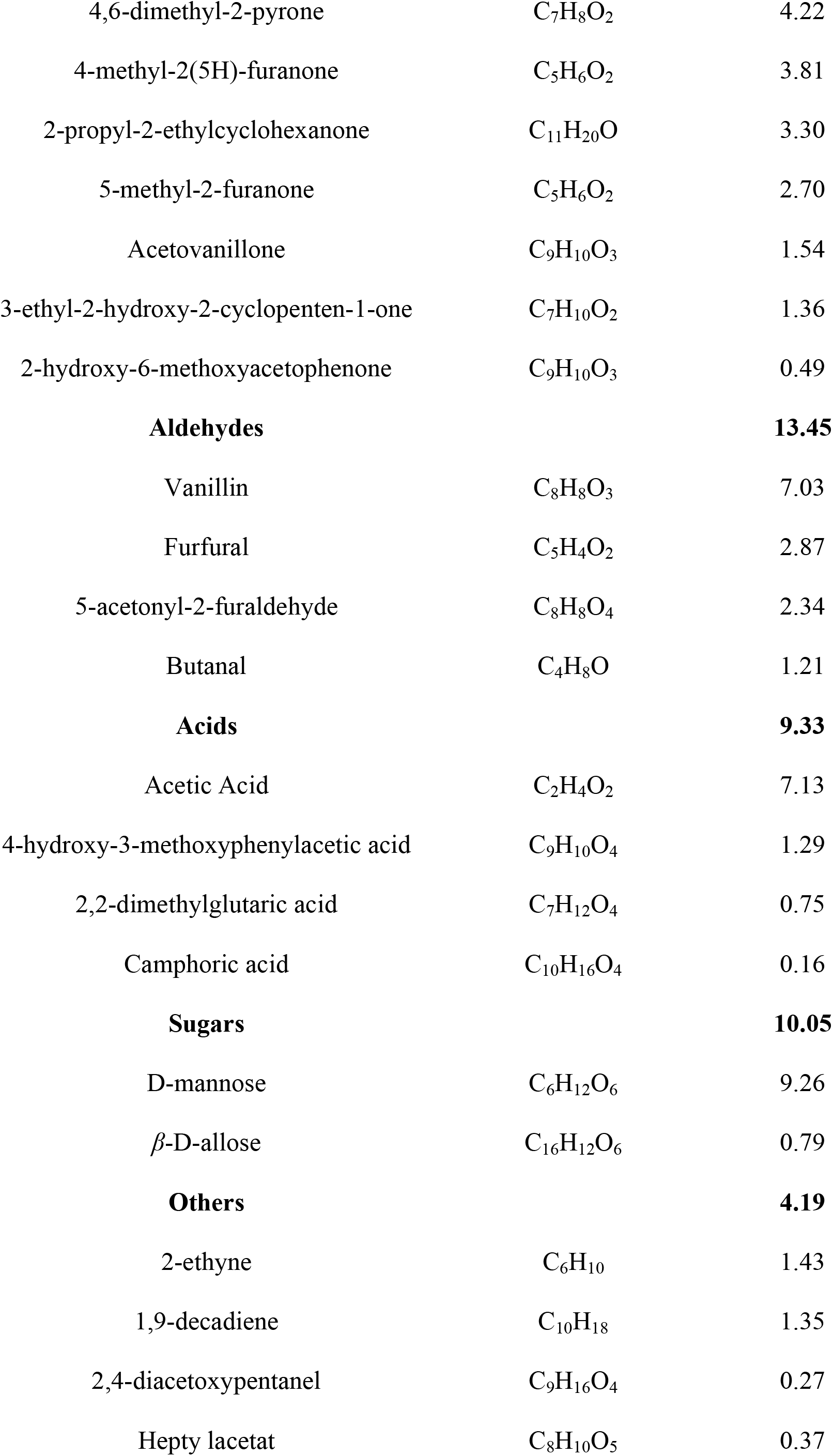
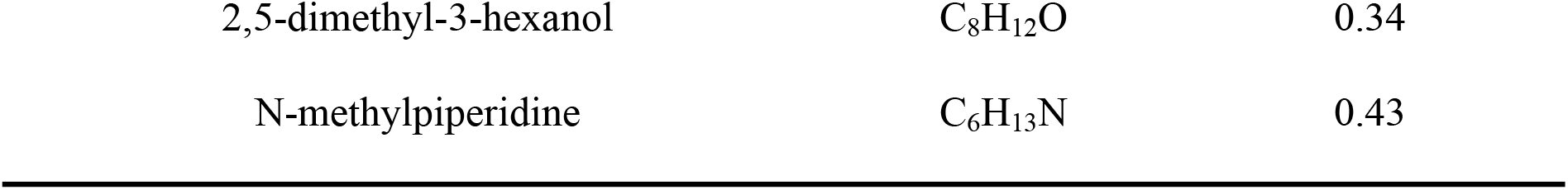
GC-MS of bio-oil.

### 2.2 Preparation

#### 2.2.1 Preparation of model compounds

According to the results of GC-MS analysis of bio-oil (Table 1), the model compounds of bio-oil were configured (Table 2). The moisture content of bio-oil was calculated as 30 %. The bio-oil and its component model compounds were prepared according to the corresponding proportions in the bio-oil. The formula and naming of bio-oil model compounds were detailed in Table 3.

**Table 2.** Composition and ratio of Bio-oil model compounds.

| Model compound | Composition | Ratio |
| --- | --- | --- |
| Phenols | Phenol; guaiacol; catechol | 4:1:2 |
| Ketones | Butanone; cyclopentane dione | 1:1 |
| Aldehydes | Vanillin; furfural | 7:3 |
| Sugars | D-glucose | — |
| Acids | Acetic acid; phenylacetic acid | 5:3 |

**Table 3.**
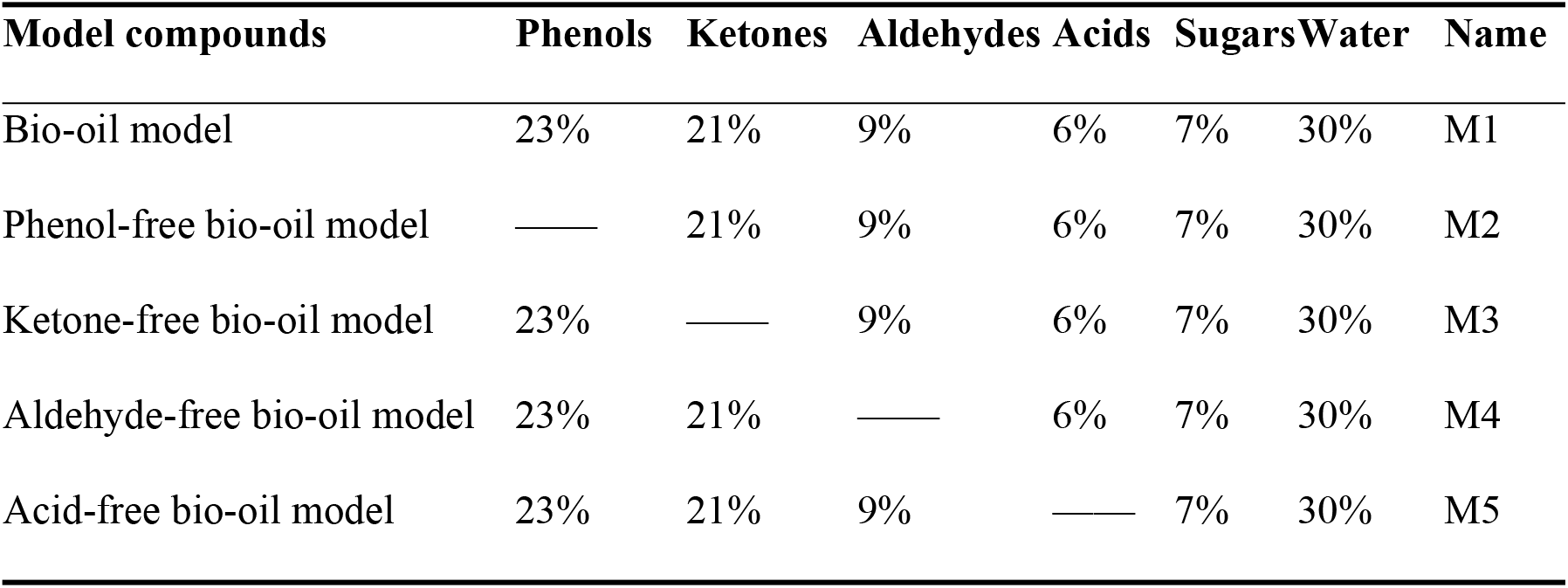
Preparation of bio-oil model compounds.

| Model compounds | Phenols | Ketones | Aldehydes | Acids | Sugars | Water | Name |
| --- | --- | --- | --- | --- | --- | --- | --- |
| Bio-oil model | 23% | 21% | 9% | 6% | 7% | 30% | M1 |
| Phenol-free bio-oil model | — | 21% | 9% | 6% | 7% | 30% | M2 |
| Ketone-free bio-oil model | 23% | — | 9% | 6% | 7% | 30% | M3 |
| Aldehyde-free bio-oil model | 23% | 21% | — | 6% | 7% | 30% | M4 |
| Acid-free bio-oil model | 23% | 21% | 9% | — | 7% | 30% | M5 |

#### 2.2.2 Preparation of resins

The BMPF were synthesized according to Yu [13]. The molar ratio of phenol (including bio-oil or bio-oil model compounds) to formaldehyde was 1:2. The substitute rate of bio-oil to phenol was 20 wt.%, and the addition amount of NaOH was 20 wt.% of the mass of phenol. The BPF with bio-oil was denoted as BPF0, and the other BPF with bio-oil model compounds were denoted as BMPF1, BMPF2, BMPF3, BMPF4 and BMPF5 according to the name of bio-oil model compounds.

#### 2.2.3 Preparation of samples

The BMPF plywood and films were prepared according to Yu [13]. After keeping for 48 h in the room, the plywood was sawed into samples with dimension of 100 mm × 25 mm × 3 mm, and 5.3 ± 0.3 g samples were selected as plywood samples. The cured resin films (80 mm × 10 mm × 2 mm) were kept in temperature humidity chamber at 20 ± 2 °C and 65 ± 5 % relative humidity for 48 h. The sample with a mass of 1.7 ± 0.2 g was selected as the resin film sample.

### 2.3 Analysis

#### 2.3.1 Basic performance test

The GC-MS analysis of bio-oil was recorded on a GC/MS-QP system (Shimadzu, Kyoto, Japan) with 20 °C/min heating rate, 3 min time-duration. An inlet temperature of 250 °C, a He gas injection volume of 0.5 μL and a flow split-ratio of 50:1 were used for the GC. A junction temperature of 260 °C, an ion temperature of 220 °C, an EI source electron energy of 70 eV and a scan range of 29-500 amu were used for MS. The moisture content of bio-oil was calculated by V30 Karl-Fischer moisture meter (Mettler Toledo, Zurich, Switzerland). The pH, viscosity, and solid content of resins were tested by China National Standards GB/T 14074-2013. Each experiment was conducted at least three times under the same conditions, and their average values were reported.

The formaldehyde emission and bonding strength of plywood was evaluated on the basis of China National Standard GB/T 17657-2013. In the test of formaldehyde emission, 10 test specimens (150mm×50 mm), cut from the plywood, were positioned in a 10 L glass desiccator. The emitted formaldehyde was absorbed by a Petri dish filled with 300 ml distilled water, and tested by the ultraviolet spectrophotometer (UV) at 412 nm wavelength. In the test of bonding strength, 12 plywood specimens with dimension of 25 mm × 10 mm × 3 mm were dipped into boiling water for 4 h, dried in 60 ± 3 °C for 20 h, and dipped into boiling water for 4 h, then submersed in cold water for 1 h. Then, the samples were tested with a speed at 10.0 mm/min of the cross head.

#### 2.3.2 Aging test

The plywood and resin film samples were examined by a UV accelerated weathering tester (Yiheng Co., Ltd., Shanghai, China) according to the ASTM G 154. Each 12 h weathering cycle consisted of 8 h of UV exposure at 60 °C and 4 h condensation at 50 °C. The plywood sample number was 6, and the resin film sample number was 3.

#### 2.3.3 Characterization

The scanning electron microscope (SEM) analysis of resin resins were measured by SU8010 SEM (Hitachi, Tokyo, Japan) at 5.0 kV accelerating voltage. The Fourier transform infrared spectroscopy (FT-IR) analysis of resin resins were tested by a Nicolet iS5 FT-IR (Nicolet, Wisconsin, USA) over the range of 400 to 4000 cm^−1^ with a 4 cm^−1^ resolution and 64 scans. The solid state nuclear magnetic resonance (NMR) analysis of resins was acquired at a frequency 100 MHz using JNM-ECZ600R (JEOL, Tokyo, Japan).

## 3 Results and discussion

### 3.1 Basic performance

The basic performance of BPF and BMPF is shown in Table 4. The pH value of BMPF were all increased in comparison with BPF0, indicating that the acidity of bio-oil model compound was lower than that of bio-oil. BMPF5 had the highest pH value because of the lack of acid compounds. Compare with BPF0, the viscosity of BMPF decreased, which might be due to the less reactions. The bio-oil model compounds had less substances compare to bio-oil, which would decrease the number of reactions. Besides, the fewer long-chain substances of bio-oil model compounds also weakened the drag effect. The low viscosity of BMPF2 was due to the lack of phenolic substances, which reduced the crosslinking degree of resin. However, BMPF3 had the lowest viscosity because the ketones were kind of good solvent and could promote the reaction between substances.

**Table 4.** Basic performance of BPF and BMPF.

| Resins | Characteristics |  |  |  |
| --- | --- | --- | --- | --- |
|  | pH<br>(25°C) | Viscosity<br>(25 °C, mPa·s) | Solid content<br>(%) | Formaldehyde<br>emission<br>(mg/L) |
| BPF0 | 11.10±0.13 | 322±54 | 45.93±0.22 | 0.272±0.021 |
| BMPF1 | 11.28±0.20 | 136±33 | 46.38±0.07 | 0.387±0.018 |
| BMPF2 | 11.33±0.07 | 101±12 | 43.40±0.10 | 0.594±0.024 |
| BMPF3 | 11.32±0.03 | 98±18 | 44.80±0.12 | 0.452±0.032 |
| BMPF4 | 11.32±0.05 | 112±25 | 46.16±0.13 | 0.539±0.039 |
| BMPF5 | 11.40±0.04 | 139±34 | 46.68±0.30 | 0.438±0.028 |

As shown in Table 4, the solid content of BMPF1 was higher than that of BPF0, indicating that the bio-oil had a greater negative effect than the bio-oil model compounds. BMPF2 had the smallest solid content, because the phenols in M2 were reduced resulting in the lower polymerization degree. The solid content of BMPF3 was smaller than that of BMPF1, which meant that ketones had a positive effect on the resin polyreaction. The increase in solid content of BMPF4 and BMPF5 indicated that the aldehydes and acids of bio-oil had a negative effect. The acids in bio-oil cloud fall the pH value of system, leading to the decreasing number of multi-substituted hydroxymethyl phenol in the resin addition process, and further reducing the resin crosslinking degree. The active site of formaldehyde is 2, while the reaction activity of other aldehydes was lower than that of formaldehyde. This meant that the other aldehydes would become the terminator of reaction, thus reducing the resin solid content.

It can be seen from Table 4 that the formaldehyde emission of BMPF1 was higher than that of BPF0, indicating that many substances in bio-oil were conducive to reduce the formaldehyde emission. In other words, there were many substances in bio-oil that could react with or adsorb formaldehyde. Besides, the formaldehyde emission of BMPF2-5 was also higher than that of BPF0, indicating that the different components of bio-oil played different positive effect on reducing the formaldehyde emission. Such as phenols can polymerized with formaldehyde, acids can be esterified with formaldehyde, and ketones can promote the reaction of other substances with formaldehyde.

### 3.2 Bonding strength

The bonding strength and its loss rate of BPF and BMPF plywood prepared are shown in Figure 1 [18]. The bonding strength of BMPF1 plywood was lower than that of BPF0 plywood, indicating that the bio-oil had less influence on the resin system than the bio-oil model compounds. BMPF2 plywood had the lowest bonding strength because the core of PF resin was formed by phenol and formaldehyde, but the total amount of phenolic substances of M2 was the lowest, resulting in a serious decrease of resin polymerization degree. Compared with BMPF1 plywood, BMPF3 plywood had a lower bonding strength, indicating that ketones could indeed promote the resin reaction. The increased bonding strength of BMPF4 resin plywood meant that the aldehydes in the bio-oil had a negative effect on the bonding strength. This might be due to 1) the aldehydes in bio-oil will compete with formaldehyde for reactive sites; 2) The aldehydes in bio-oil basically have only one active site, which blocked the further increase of resin molecular chain. Compared with BMPF1 plywood, the bonding strength of BMPF5 plywood increased, which was due to the decrease of pH value.

**Fig 1.**
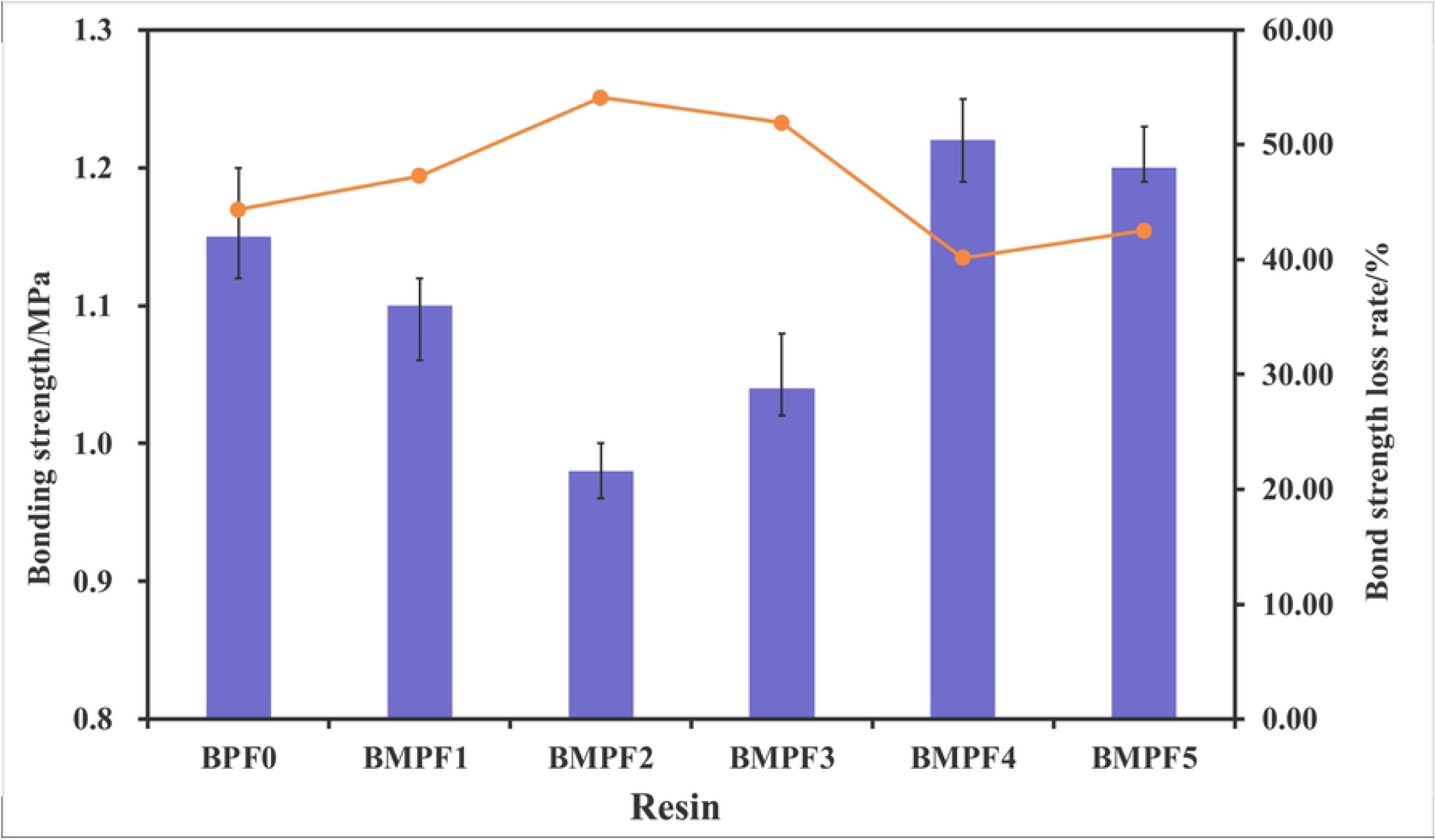
Bonding strength and its loss rate of BPF and BMPF plywood.

It can be seen from Figure 1 that after 960 h of aging, the loss rate of bonding strength of BMPF1 plywood was higher than that of BPF0 plywood, indicating that except the bio-oil model compounds, the other substances in bio-oil also played a positive role in aging resistance. The biggest bonding strength loss rate of BMPF2 plywood was due to the decrease of phenolic content. Compared with BMPF1 plywood, the bonding strength loss rate of BMPF3 plywood was higher, which was due to the long-chain flexible groups in ketones. As all known, the long-chain flexible groups could effectively improve the toughness of resin. Besides, the increased toughness would reduce the resin layer fractures caused by the stress generated and water-corrosion during aging, thereby improving the aging resistance of resin. The decrease of bonding strength loss rate of BMPF4 and BMPF5 plywood was due to the negative effect from aldehydes and pH value on resin synthesis reaction. However, the bonding strength loss rate of BMPF5 plywood was higher than that of BMPF4 plywood, because the lower pH value cloud reduced the resin solubility, which slightly improved the water resistance.

### 3.3 SEM analysis

The apparent morphology changes of BPF and BMPF films before and after aging are shown in Figure 2 [14,19]. The BMPF films before aging had a smooth surface, and the degree of smoothness was higher than that of BPF0. The reason was that there were some solid particles in the bio-oil, which reduced the smoothness of its solidified surface. Besides, it also showed that the selection of bio-oil model was reasonable, which produced better synthesis reaction and formed a smooth surface after curing. Compared with BMPF2-5, the smoothness of BMPF3 film surface was lower because ketones can act as a good solvent and promote the resin to form a better surface.

**Fig 2.**
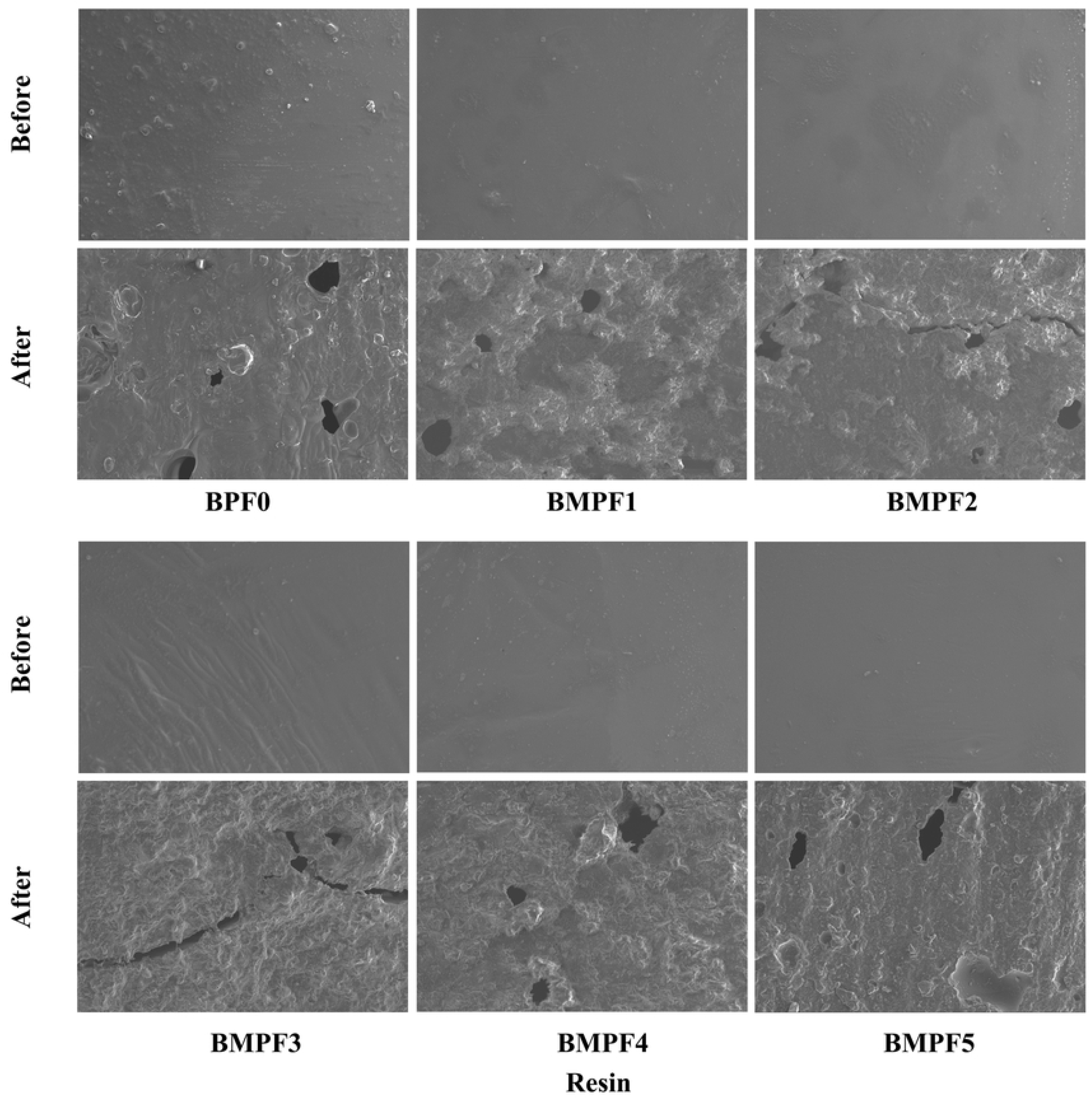
SEM images of BPF and BMPF films before and after aging for 960 h (× 500).

It can be seen from Figure 2 that the surface roughness of BPF and BMPF increased obviously after aging for 960 h, and defects such as holes and cracks appeared. Compared with BPF0, BMPF1 film had a higher degree of surface aging. Compared with BMPF1-5, BMPF3 and BMPF2 had a greater aging degree. The reduction of phenolic substances reduced the degree of resin polymerization, thereby weakening the aging resistance. Meanwhile, the lack of ketones reduced the oily substances in the system, resulting in a lower water resistance and a higher degree of surface aging of BMPF2 and BMPF3.

### 3.4 FT-IR analysis

The FT-IR curves of BPF and BMPF before and after aging for 960 h are displayed in Figure 3, and the attribution of characteristic absorption peaks is shown in Table 5 [12,13,15,29]. As shown in Figure 3, the peak of methylene (CH_2_) groups at the region of 2924 cm^−1^ and 2854 cm^−1^. The content of CH_2_ groups can indirectly represent the polymerization degree of PF resin [30,31]. The CH2 groups peak of BMPF1 was stronger than that of BPF0, because the phenolic substance activity of bio-oil model was stronger than that of bio-oil. Compared with the BMPF1, the CH_2_ groups peak of BMPF2 was weaker because CH_2_ was generated by the reaction of phenols and aldehydes, and the reduction of phenols reduced the CH_2_ content in the resin. The weakening of CH_2_ groups peak of BMPF3 also confirmed the promoting effect of ketones on resin polymerization. The increased of CH_2_ groups peak of BMPF4 and BMPF5 meant that aldehydes and acids in bio-oil have a negative effect on the synthesis of BPF resin. The peak of ether bond (C—O—C) groups at the region of 2924 cm^−1^ and 2854 cm^−1^. Ether bond is another connecting structure between molecular chains of PF resin, which can also indirectly represent the degree of polymerization of PF resin. Compared with BPF0, the change on ether bond peak of BMPF1-5 was similar to those of CH_2_ peak.

**Fig 3.**
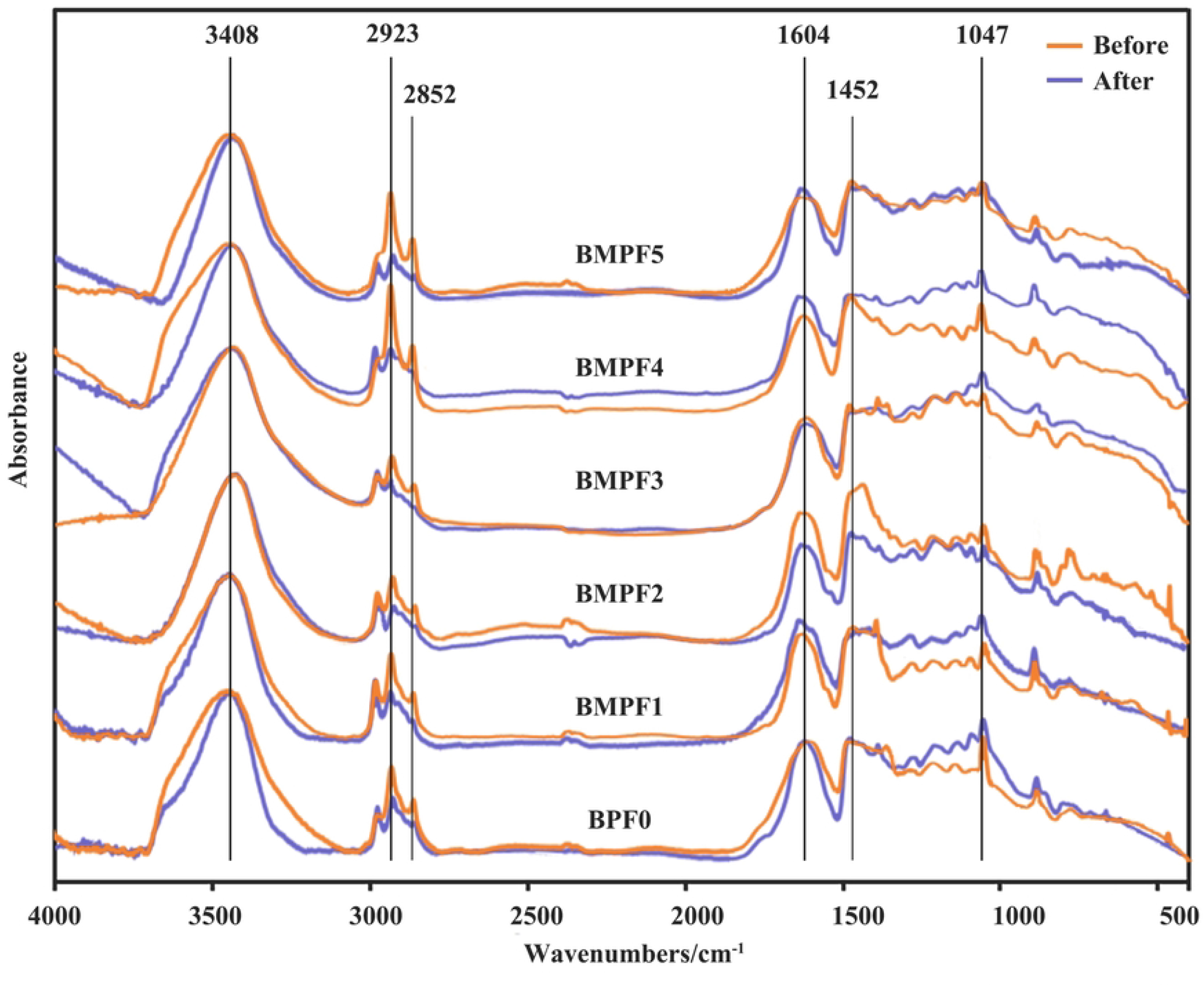
FTIR curves of BPF and BMPF before and after aging for 960 h.

**Table 5.** Peaks and assignment of FT-IR spectra for BPF.

| Wave number (cm<sup>-1</sup>) | Vibration | Assignment |
| --- | --- | --- |
| 3408 | $\nu(\text{—OH})^a$ | Phenolic OH and aliphatic OH stretching vibration |
| 2923, 2852 | $\nu(\text{CH}_2)$ | Aliphatic CH<sub>2</sub> asymmetric stretching vibration |
| 1604, 1452 | $\nu(\text{C}=\text{C})$ | C=C aromatic ring stretching vibration |
| 1047 | $\nu(\text{C—O—C})$ | Phenolic C—O—C stretching vibration |

<sup>a</sup> $\nu$ : Stretching vibration.

After aging for 960 h, the peak of CH_2_ and C—O—C groups decreased significantly, indicating that the polymerization degree of resin decreased after aging. Compared with BPF0, the CH_2_ peak of BMPF1 decreased significantly. It showed that other substances in bio-oil could reduce the polymerization degree of resin synthesis, but improved the aging property of resin. Compared with BMPF1, the CH_2_ groups peak of BMPF2 decreased significantly after aging, which might because resin failed to form a good network structure due to the reduction of phenols, resulting in the reduction of aging resistance. The CH_2_ groups peak of BMPF3 weakened the biggest. It might be due to the lack of ketones, which increased the contact area of external environment (water, light and other aging factors), and aggravated the aging degree.

### 3.5 NMR analysis

^13^C NMR images of BPF before and after aging for 960 h are displayed in Figure 4. The chemical shifts of carbon atoms in each group of BPF resin are shown in Table 6 [18,32]. The carbon spectra of BMPF1-5 and BPF0 were basically similar, indicating that the preparation of bio-oil model compound was reasonable, and the conclusion was the same as that of FT-IR analysis. BPF0 had more substances with C=O structure than BMPF1-5 because there were more kinds of C=O substances in bio-oil. Compared with BMPF1, the values of substituted ortho and para aromatic carbons (139-120 ppm) and methylene bridges (45-27 ppm) of BPF0 were smaller, indicating that other components of bio-oil would still affect the resin synthesis process. However, the value of dimethylene ether bridges (77-68 ppm) increased, which might be due to the presence of ethers in bio-oil. Compared with BMPF1, the values of methylene bridges of BMPF2 and BMPF3 decreased, indicating that phenols and ketones in bio-oil played a positive role in synthesis reaction, while the values of methylene bridges of BMPF4 and BMPF5 increased, indicating that aldehydes and acids in bio-oil had opposite effects.

**Fig 4.**
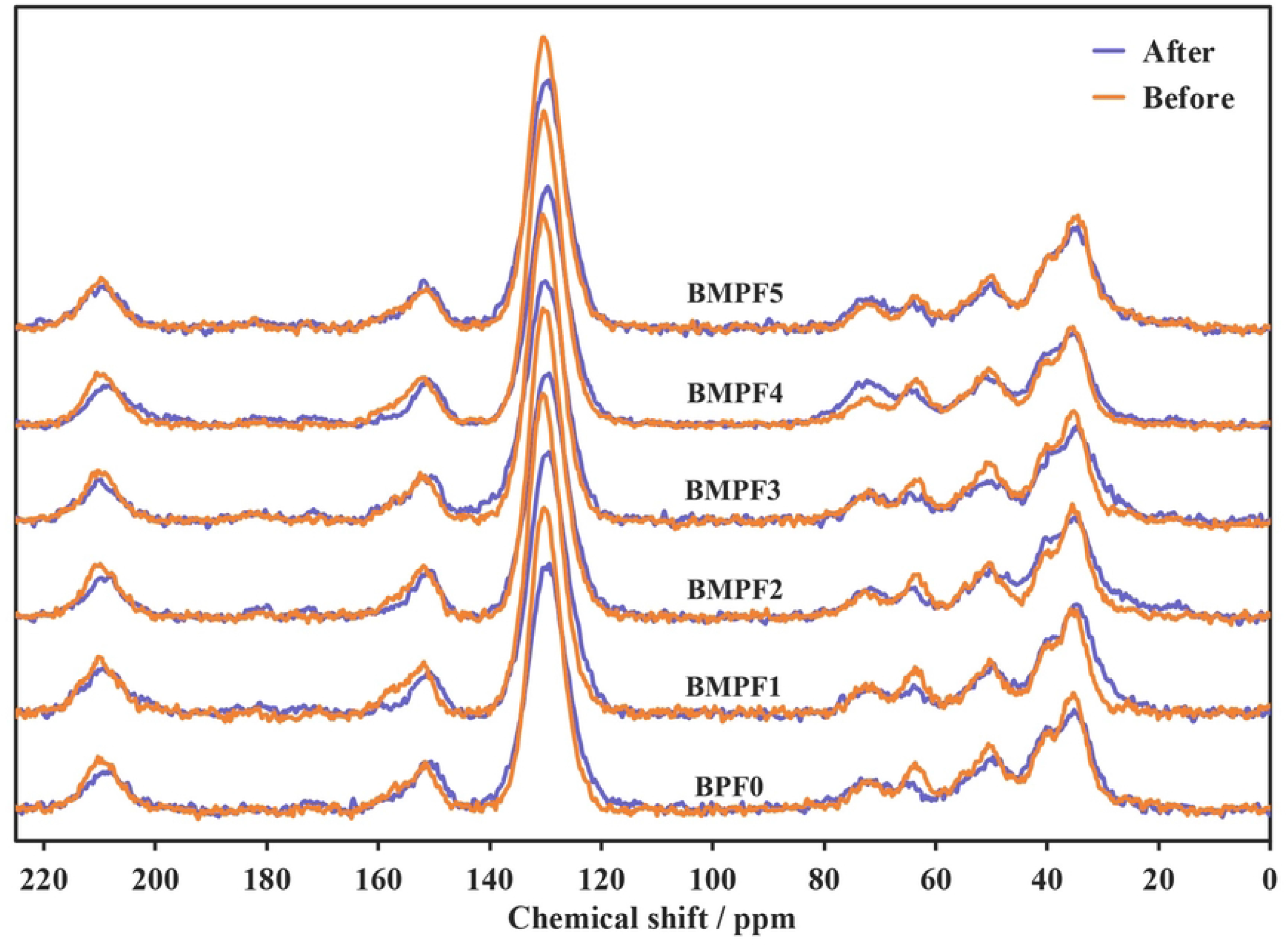
^13^C NMR images of BPF and BMPF before and after aging for 960 h.

**Table 6.**
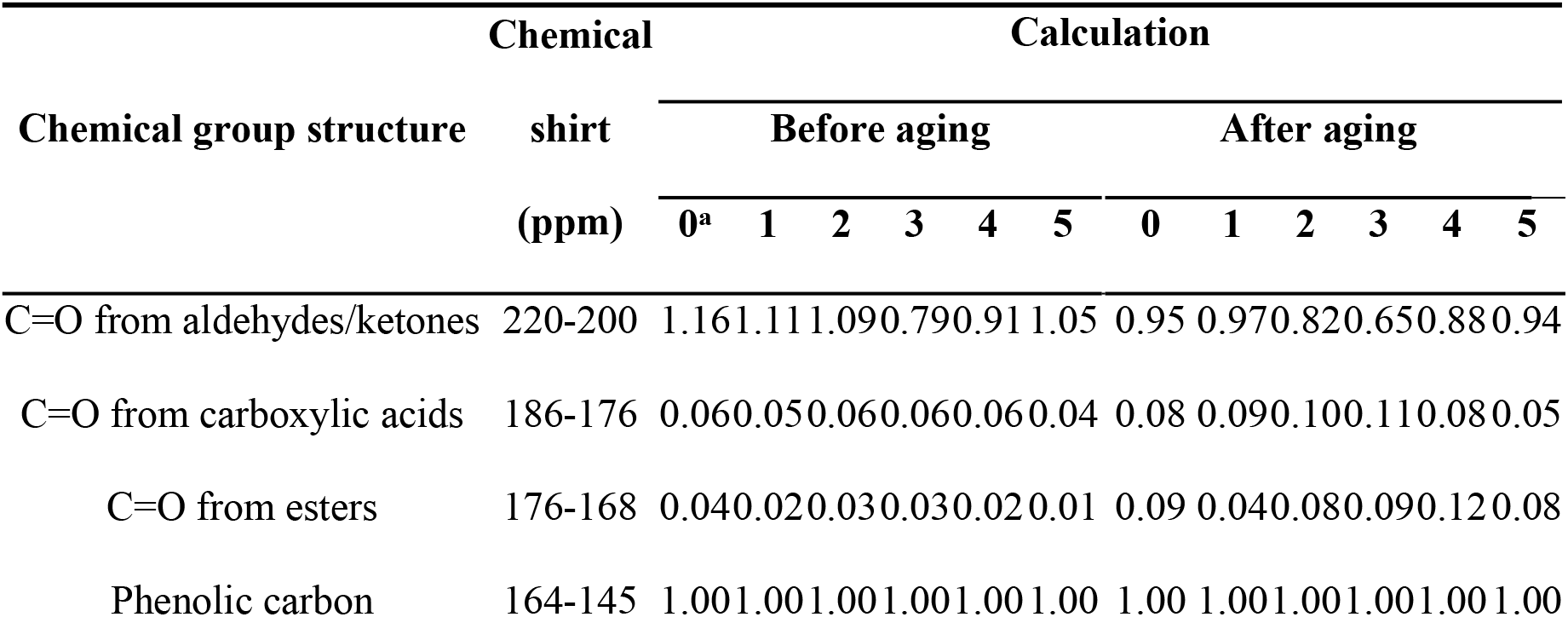
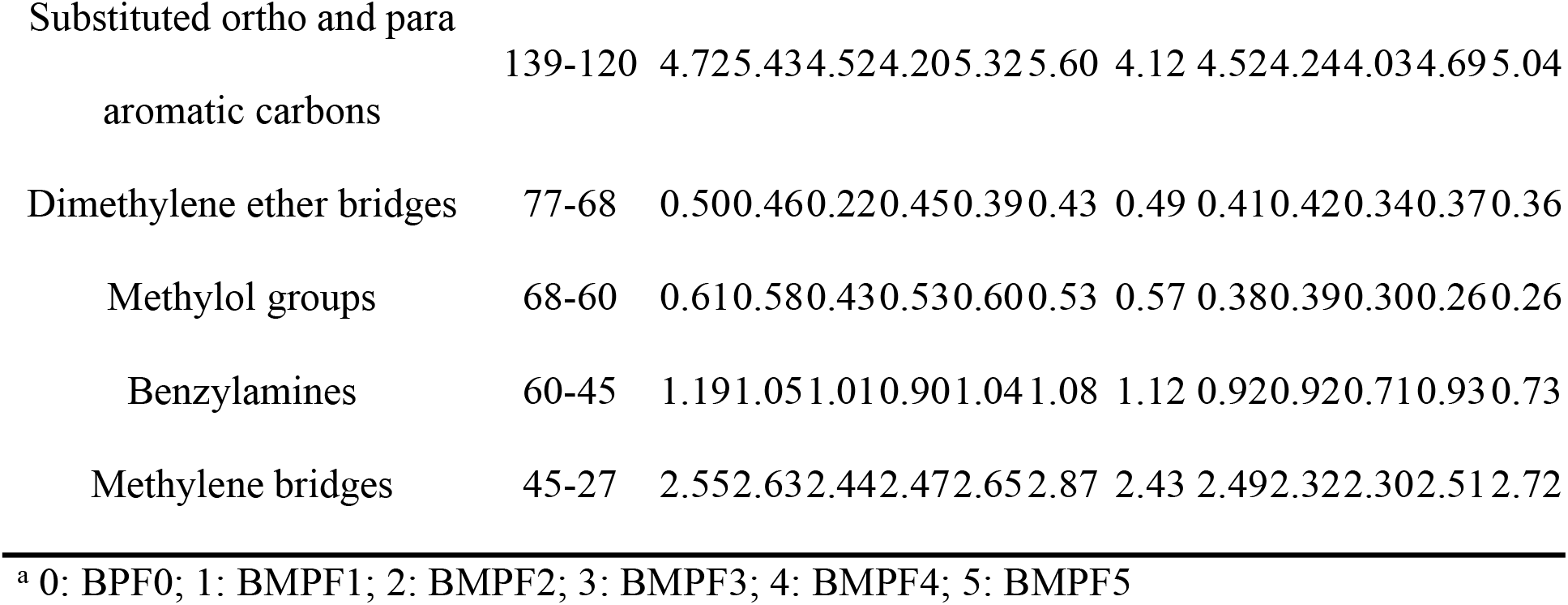
^13^C NMR assignment and quantitative analysis of chemical groups for BPF before and after aging for 960 h.

After aging for 960 h, the intensity of C=O from carboxylic acids and esters (186-168 ppm) increased, indicating that the aged resin had undergone oxidative aging. The intensity of C=O from aldehydes (220-200 ppm) decreased, indicating that some aldehyde groups were converted to carboxyl or ester groups. The reduction of the substituted ortho and para aromatic carbons meant that the resin network structure was broken after aging, and small molecular substances were formed. The decrease of the intensity of methylene bridges indicated that the molecular chain of resin was broken after aging. Compared with BMPF1, the reduction of the methylene bridge of BPF0 was lower, indicating that the improvement on aging resistance of BPF was the result of the overall interaction of bio-oil molecules. The reduction of methylene bridge of BMPF2-5 was higher than that of BMPF1, which also confirmed this inference.

## Conclusions

Crude bio-oil was used as a phenol substitution to synthesize the BPF resin, and its performance and modification mechanism were analyzed by model compound method. The bio-oil components had different influence on performance and microstructure of BPF. Compared with BMPF1, the bonding strength of BMPF2 and BMPF3 plywood decreased, while the bonding strength of BMPF4 and BMPF5 increased. Structural analysis showed that the CH_2_ peak in FT-IR and the methylene bridges intensity in NMR of BMPF2 and BMPF3 were lower than that of BMPF1, while the results for BMPF4 and BMPF5 were opposite. These indicated that the phenols and ketones of bio-oil had positive effects, while the aldehydes and acids had negative effects. After aging for 960 h, compared with BMPF1, the CH_2_ peak of BMPF2-5 decreased significantly and the reduction of methylene bridge from BMPF2-5 was higher, which meant that all components of bio-oil could improve the aging resistance of BMPF inordinately. These results could provide a basis for the modification of BPF.

## Acknowledgements

This work was supported by the Talent Introduction Program of Zhejiang Sci-Tech University (19082425-Y) and Philosophy and Social Science Planning Project of Zhejiang Province (21NDQN234YB).

